# Inherent Biomechanical Properties of the Lung: *In vivo-Ex vivo* Comparisons in Mice

**DOI:** 10.64898/2026.06.24.734270

**Authors:** Jack Di Palo, Jonathan T. Ibinson, Liqin Lin, Bryan Suh, Meredith S. Gwin, Sandra Zaeh, Jason M. Szafron, Edward P. Manning

**Affiliations:** Section of Pulmonary, Critical Care and Sleep Medicine, Yale School of Medicine, Yale University, New Haven, CT, US; Department of Biomedical Engineering, Carnegie Mellon University, Pittsburgh, PA, US

## Abstract

Mammalian lungs operate within a thoracic “cage” composed of parietal pleura, rib cage, skeletal muscle, and diaphragm, yet clinical ventilator metrics largely reflect the combined mechanics of lung and surrounding structures and the thoracic cage. We hypothesized that thoracic boundary conditions selectively alter measured lung biomechanics. We performed paired pulmonary function testing (FlexiVent) in C57BL6 mice of both sexes spanning development through adulthood, measuring quasi-static pressure–volume behavior and dynamic forced-oscillation parameters *in vivo* (supine, mechanically ventilated) and again *ex vivo* in the same lungs. In a subset, we additionally compared *in vivo* and *ex vivo* µCT-derived lung volumes, including a pressure-fixed *ex vivo* protocol using snap freezing at controlled inflation pressure. Quasi-static pressure–volume curves were similar between conditions, with near-identity at higher pressures and only modest divergence at low pressures, consistent with thoracic structures primarily modulating recruitment/de-recruitment rather than intrinsic elastic recoil. Maximal volume at 30 cmH2O showed strong *in vivo-ex vivo* correlation and minimal bias, and static compliance and PV-loop hysteresis exhibited small biases relative to reported disease-model effect sizes. In contrast, dynamic mechanics demonstrated a clear *in vivo* elevation of tissue damping (G) with only modest change in tissue elastance (H) and little change in Newtonian resistance (Rn), producing a meaningful increase in hysteresivity (η = G/H). This dissociation implicates frequency-dependent mechanical heterogeneity (e.g., time-constant mismatch/pendelluft) imposed or amplified by nonuniform thoracic loading. *Ex vivo* µCT enabled reliable whole-lung segmentation and correlated with *ex vivo* PFT volumes at matched pressures, whereas *in vivo* volumetry showed weaker agreement. These results indicate that thoracic structures contribute modest restriction but disproportionately increase dynamic dissipation and heterogeneity, suggesting that *ex vivo* functional testing and oscillometry-like metrics may better detect biomechanical changes inherent to lung parenchyma.

## INTRODUCTION

Mammalian lungs are confined within the thoracic cavity. As such, lung parenchyma and airways are surrounded by pleura, bones, skeletal muscle, and diaphragm with several openings for the trachea, esophagus, veins, arteries, and nerves. It is reasonable to assume that the thoracic cavity’s boundaries act as a “cage” that counters expansion of the lungs during inflation, prevents deflation or atelectasis, and impact lung mechanics.

Delineating mechanical properties of the lung is clinically important. Increased understanding of lung airway and parenchymal mechanics has advanced safe and effective use of mechanical ventilation (1, 2). This has improved our ability to provide therapeutic supplemental oxygen for patients with hypoxemic respiratory failure such as pneumonia and acute respiratory distress syndrome (ARDS), while preventing ventilator-induced lung injury (VILI) (1, 2). With ARDS patients specifically, significantly improved mortality rates have been shown when we understand and control the mechanical properties of lung tissue (3, 4).

However, understanding of the biomechanical properties of lung parenchyma remains limited partially due to the lung’s interaction with the thoracic cavity. For example, mechanical ventilation uses the biomechanical metrics of positive end-expiratory pressure (PEEP) and plateau pressure to properly set ventilation parameters (5). However, these measurements reflect properties of the entire respiratory system and fail to discern properties inherent to the lung from those of surrounding structures. Some current techniques, like esophageal manometry aim to better identify the lung mechanical properties by quantifying transpulmonary pressure. This pressure measurement allows for the separation of lung parenchyma and airway mechanics from the influence of surrounding structures (1, 2). However, quantifying transpulmonary pressure is not currently common in clinical practice (6). Additionally, it is challenging for pulmonologists and respiratory therapists to manage patients with parenchymal lung disease when the patient also suffers from abnormal chest physiology or injury. For example, a patient with pulmonary fibrosis may also have flail chest. These counteracting factors may result in normal biomechanical values of lung function, such as normal plateau pressures during mechanical ventilation, despite having underlying lung disease where abnormal mechanical values are expected (1, 2). Finally, the patient anatomy itself can be a factor, as chest wall size has also been shown to abnormally contribute to the ventilation parameters in a clinical setting (7). The combination of these factors can distort the ventilation parameters on which physicians base their clinical decisions.

We hypothesized that thoracic components (bone, muscle, etc.) affect the biomechanical properties of the lung. To test this hypothesis, we performed pulmonary function tests (PFTs) and imaging analysis on *in vivo* and *ex vivo* micro-computed tomography (*u*CT) of male and female mouse lungs of varying ages throughout development until adulthood.

## METHODS

### Mice

The study was approved by the Yale University Institutional Animal Care and Use Committee. C57BL/6J mice of both sexes ranging in age from 3 to 18 weeks and 27 months old (normative development and aging) were housed in an antigen-free and virus-free animal care facility under a 12-h light and dark cycle. Animals were euthanized in accordance with approved IACUC protocols. Thoracic diameter measurements of these mice were included in our previous publication (8).

### Pulmonary Function Testing

Subjects were anesthetized via an intraperitoneal injection of urethane (1 g/kg of body weight in 10% solution with sterile water). They were subsequently tracheostomized and connected to the FlexiVent system (FlexiVent, SCIREQ, Montreal, QC, Canada). Following connection to the FlexiVent, an intraperitoneal injection of pancuronium (1 mg/kg) was administered to passivate the subject’s breathing. Four standard FlexiVent maneuvers were performed according to the manufacturer: Deep Inflation, a pressure-driven maneuver to inflate the lungs up to total lung capacity at 30 cmH_2_O (reported as inspiratory capacity, IC); Snapshot Perturbation, a single-frequency forced oscillation maneuver to measure global lung mechanics as a function of pressure, flow, and volume changes resulting from that oscillation (reporting respiratory system elastance (Ers) and dynamic compliance (Crs)); Quick Prime Perturbation, a multi-frequency forced oscillation maneuver to measure frequency-dependent lung mechanics distinguishing airway versus parenchymal mechanical properties of the lungs (reported as airway resistance (Rn), tissue damping (G), parenchymal elastance (H), and hysteresivity (η, the ratio of G/H); and a Pressure-Volume loop, a quasi-static stepwise inflation/deflation of the lungs measuring lung distensibility independent of flow (reporting static compliance (Cst), total lung capacity similar to Deep Inflation (IC), and hysteresis reflecting tissue energy storage and dissipation during recruitment and de-recruitment) (9). This block of four maneuvers was repeated three times per subject per condition (*in vivo* vs *ex vivo*). The raw data was analyzed using the FlexiWare Version 7.6 software, Service Pack 6 to obtain pulmonary function parameters described above. For each parameter, only measurements meeting the system’s coefficient of determination (COD) quality threshold were retained, and a parameter was accepted for an animal only when three consistent, non-flagged runs were obtained. Animals failing this criterion for a given maneuver were excluded from that parameter. Because different parameters are derived from different maneuvers, the total number of animals contributing to a reported parameter varies.

The *in vivo* PFT was performed with the mouse in a supine position (Figure 1A). Immediately after the *in vivo* PFT, the mice were euthanized in accordance with our IACUC-approved protocols, and the heart and lungs were removed *en bloc*. The trachea was cannulated and the PFT (the same four maneuvers performed *in vivo*) was repeated *ex vivo*, with the heart-lung block resting in a dish filled with phosphate buffered saline (PBS). Following the *ex vivo* PFT, the lungs were snap frozen with liquid nitrogen at 20 cmH_2_O of pressure via a gravity-fed fluid column of PBS. The *ex vivo* PFT shows both pressure extremes from 3 cmH_2_O (Figure 1B) to 30 cmH_2_O (Figure 1C).

**Figure 1.**
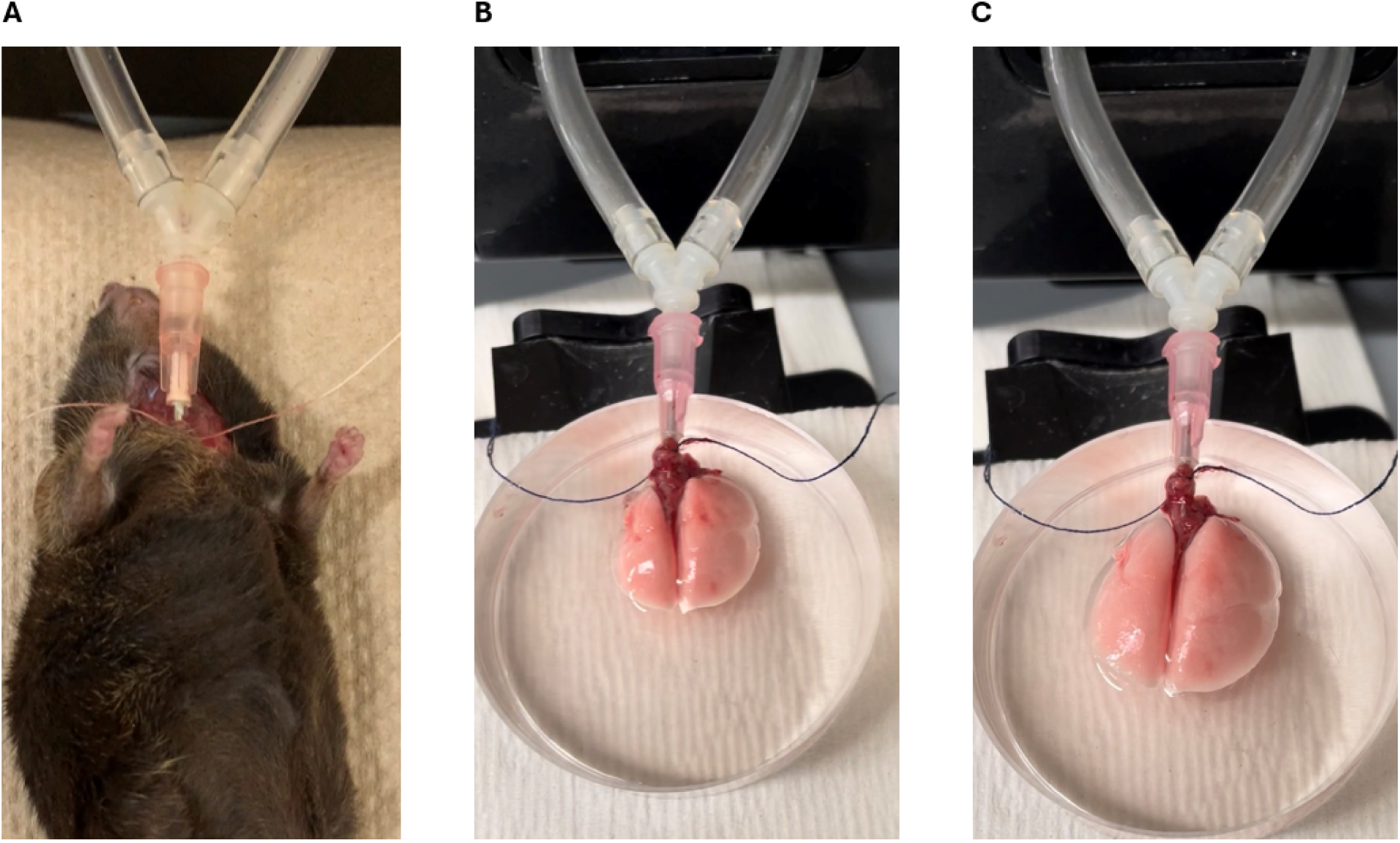
*In vivo* and *ex vivo* Pulmonary function test (PFT). A) *In vivo* PFT B) *Ex vivo* PFT at the beginning of the deep inflation maneuver at 3 cmH_2_O of pressure C) *Ex vivo* PFT at the end of the deep inflation maneuver at 30 cmH_2_O

### *In vivo u*CT imaging

*In vivo* imaging was done prior to the PFT. Each subject was sedated with 1.5% isoflurane delivered via nose cone. Each was scanned in a supine position using a SPECT/high-resolution CT scanner (MI Labs USPECT4/CT). A motion sensor was strapped to the subject’s chest to track thoracic excursion during respiration. This allowed for segmentation of the respiratory cycle into four phases from maximum inspiration to expiration. Images were sorted according to phase in the respiratory cycle during post-processing.

From imaging, we quantified volumes under load. To conduct this volumetric analysis, the total lung volumes from the above images were segmented in 3D Slicer (10). Our semi-automated segmentation process began with the open-source extension Lung CT Analyzer (11, 12), which generated rough volumes of the lung anatomical features from user inputted seed points. Due to chest compression from the motion sensor probe, the left and accessory lung lobes were not visible in most samples (Figure 2A). Thus, only the right cranial, middle, and caudal lobes were analyzed for the *in vivo* samples. Once volume estimates were formed through the Lung CT Analyzer algorithm, segmentations were manually smoothed and filled to better match each subject’s lung position. *In vivo* lung volumes were then extracted from the generated lung mask (Figure 2B). Finally, the thoracic lung area was measured in an axial 2D slice of the thoracic cavity directly above the diaphragm, as an easily determined whole lung-size metric that was independent of the lost left/accessory lung volumes (Figure 2C).

**Figure 2.**
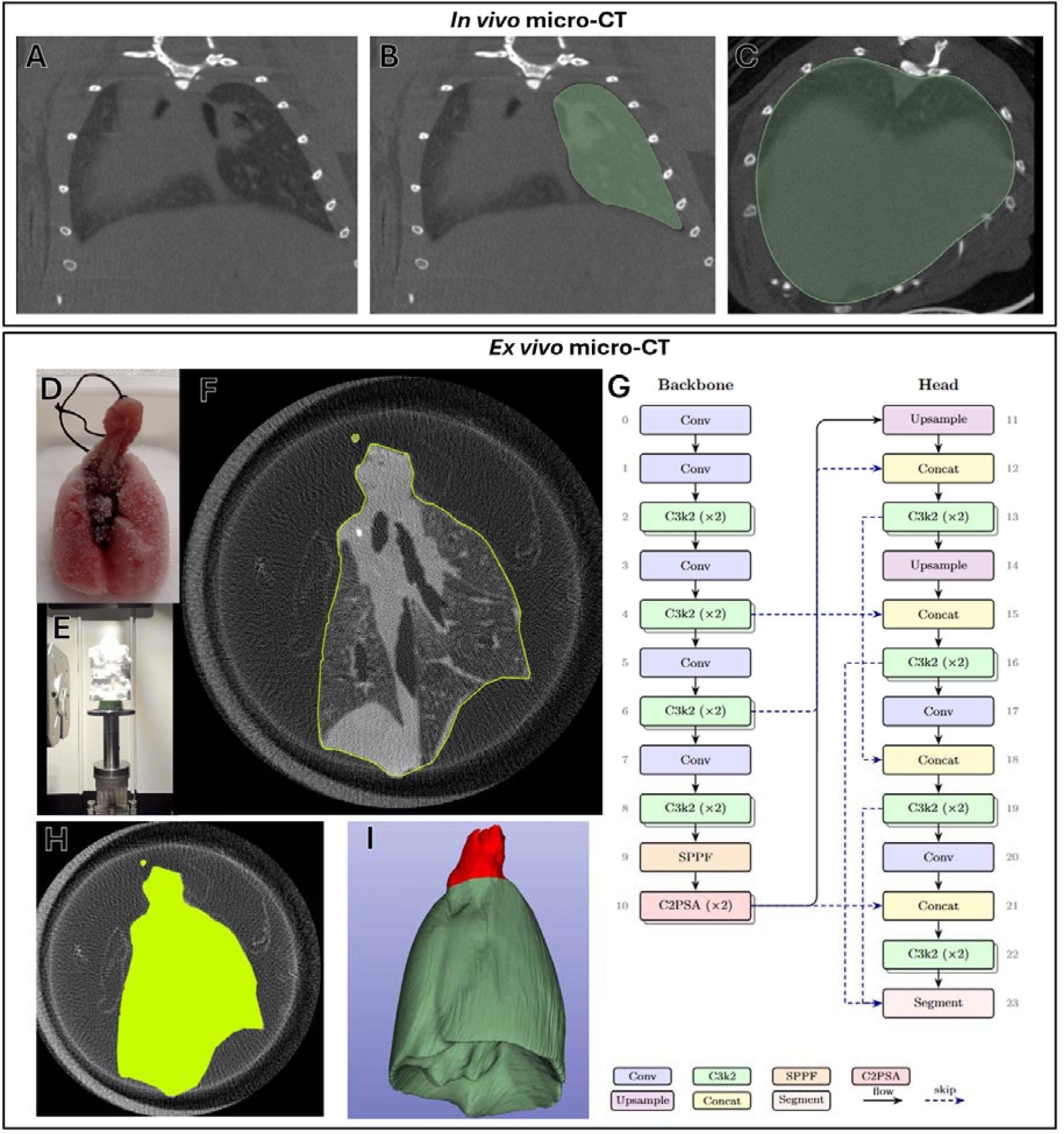
**A–B)** Example *in vivo* images before and after segmentation illustrate that *in vivo* segmentation had difficulty identifying aerated volumes in the left lung lobes. **C)** To address this, one feasible alternative is to estimate total lung volume from *in vivo* µCT by extracting and summing 2D axial lung-field areas. **D)** *Ex vivo* lungs can be inflated to defined pressures and fixed in that state by snap-freezing over liquid nitrogen. **E)** The frozen, pressure-fixed lung can be maintained in its conformation during scanning by placing it in a temperature-controlled Falcon tube surrounded by insulation on the µCT stage within the scanner. **F-G)** This approach produces high-quality µCT images of frozen lungs at a target pressure, which can then be manually annotated or automatically segmented using our algorithm (see Methods) to generate fully segmented lung volumes. **H-I)** The workflow enables whole-lung segmentation; the lung volume is retained (green) while the trachea is excluded (red)

### *Ex vivo u*CT imaging

After dissection, the frozen lungs were stored and shipped to Carnegie Mellon University on dry ice (Figure 2D). A total of 8 frozen lung samples (5 male, 3 female) were imaged *ex vivo* and analyzed to calculate total lung volumes. The inflated, frozen lungs were scanned using a Zeiss Xradia CrystalCT (Oberkochen, Germany) system. Samples were scanned for an exposure time of 2 seconds for 401 projections with a target voltage of 80kV, a target power of 7W, and a resolution of approximately 10μm. To ensure the mouse lungs remained frozen for the duration of the scan, the lungs were placed in a 50 mL Falcon tube (Tewksbury, MA, USA) with pebbled dry ice surrounding the sample. The sample was encased in a Kimtech Science Kimwipe (Roswell, GA, USA) to separate the lungs from direct contact with the dry ice. Minimal dry ice was placed below the sample to avoid motion artifacts as the ice melts. The tube was surrounded with insulating foam rings to reduce heat transfer, and a hole was punctured at the top of the Falcon tube to allow for CO_2_ dissipation (Figure 2E). The resulting scan was reconstructed using the Zeiss Xradia Scout-and-Scan Reconstruction version 16.2 (Oberkochen, Germany) software.

Lung volumes were segmented from the reconstructed stack using a convolutional neural network-based pipeline, featuring a fine-tuned YOLO11s-seg model (13). An automated pipeline was used to analyze these images (Figure 2F-I).

To prepare the reconstructed images for post-processing, the raw μCT volumes were reduced from approximately 3000×3000×1930 to 1000×1000×643 using voxel binning in FIJI (14). This process suppresses noise for human annotators, making the lung area easier to label slice-by-slice in Roboflow (15) using Segment Anything Model 3 (SAM 3)(16). These labels were then exported with domain appropriate augmentations and were used to train a series of YOLO11s-seg models via the Ultralytics framework (https://github.com/ultralytics/ultralytics) using four NVIDIA H100s provided by Carnegie Mellon University’s ORCHARD cluster. Training proceeded in three stages with mask ratio values of 4, 2, and 1; the first model was trained from randomized weights for 200 epochs with the MuSGD optimizer (17), an image size of 1024px, and a patience value of 50. Each subsequent run was initialized from the previous run’s weights using the same hyperparameters. This staged reduction ensured training stability compared to initializing with a mask ratio of 1 while producing the highest-quality masks. Manual annotations (Figure 2F) were used in place of model predictions for the volumetric results whenever available.

A semi-automated pipeline was used to apply significant post-processing to the raw output of the model, excluding one volumetric result with manual segmentation. First, the concave diaphragmatic surface of the lungs cannot be accurately represented by the simply-connected 2D output of YOLO instance segmentation in certain orientations. As such, volumes were resliced along one or more orientations to minimize the contributions of such interior holes being incorrectly filled, then projected back into the native coordinate space, and combined by voxel-wise union. Second, during prediction, the Intersection over Union (IoU) threshold for Non-Maximal Suppression (NMS) was set to 1 to retain all candidate masks. However, this yields a foreground with interior holes when overlapping masks enclose a low-confidence region. In contrast to the process above, these holes are an artifact rather than a desired feature and were filled by the pipeline. Third, confidence values used during prediction were manually tuned for each sample to suppress false positive detections. The false negative detections were recovered by applying linear interpolation between the nearest foreground masks. Finally, the region of interest was manually isolated in 3D Slicer by removing the trachea and structures superior to it (Figure 2I) as well as spurious detections of extraneous tissue. The *ex vivo* lung volumes were computed by summing the remaining foreground voxels and converting to physical units.

### Statistical Methods

*In vivo* and *ex vivo* measurements were performed on the same set of lungs for each mouse. Therefore, we compared values using pairwise testing. For each reported FlexiVent PFT parameter, the difference between the *in vivo* and *ex vivo* values was computed and the distribution of this difference for each parameter was assessed for normality using the Shapiro-Wilk test. Reported p-values greater than 0.05 were treated as normally distributed. For normal distributions, paired t-tests were run on the *in vivo* and *ex vivo* values. Only PFT parameter G, tissue damping, failed the Shapiro-Wilk test for normality. For this parameter, instead of a paired t-test, we used the Wilcoxon signed-rank test to analyze differences. For both the t-test, and Wilcoxon test, a two-tailed p-value of less than 0.05 was considered statistically significant.

As biomechanical measurements are continuous in nature, we compare variances between the *in vivo* and *ex vivo* paired measurements using linear regression. We used Bland-Altman (difference) plots to assess the agreement between *in vivo* and *ex vivo* measurements. This is a useful method to determine if two observations of the same metric give similar results without clinically meaningful differences, or in pre-clinical studies relative to reported effect sizes in mouse models. This method identifies bias by comparing the mean differences between the two methods. A mean difference of zero or near zero indicates that there is no systemic bias for either method of measurement. The limits of agreement identify the range of random variation between which most measurements are found. If the value of the limits are not greater than physiologically or clinically meaningful values, the two methods are considered equivocal for experimental or clinical use, respectively.

All statistics were run in GraphPad Prism version 11.0.1.

## RESULTS

We performed PFTs on 21 C57BL/6 mice (9 female, 12 male), of ages ranging from 3 to 18 weeks and an additional group of age 27 months. Data on these mice are provided in Table 1. Representative images of *in vivo* and *ex vivo* PFTs are shown in Figure 1. Videos of *ex vivo* PFTs are provided in Supplemental Information.

**Table 1.**
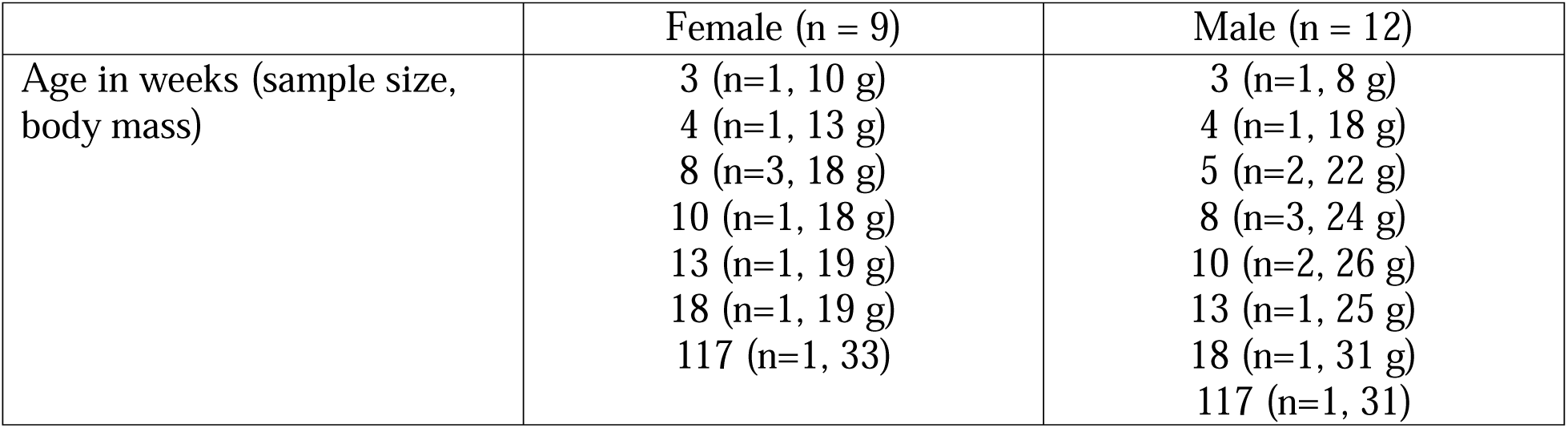

First, we compared *in vivo* and *ex vivo* results for static and quasi-static parameters. The pressure-volume relationship of *in vivo* and *ex vivo* lungs is similar (Figure 3A). When the deflation limb of the PV curves is above 15 cmH_2_O, the loops are nearly identical. Comparatively, there is a modest upward shift of the *ex vivo* loop at the lower pressures of the deflation limb (less than 15 cmH_2_O) and throughout the entirety of the inflation limb.

**Figure 3.**
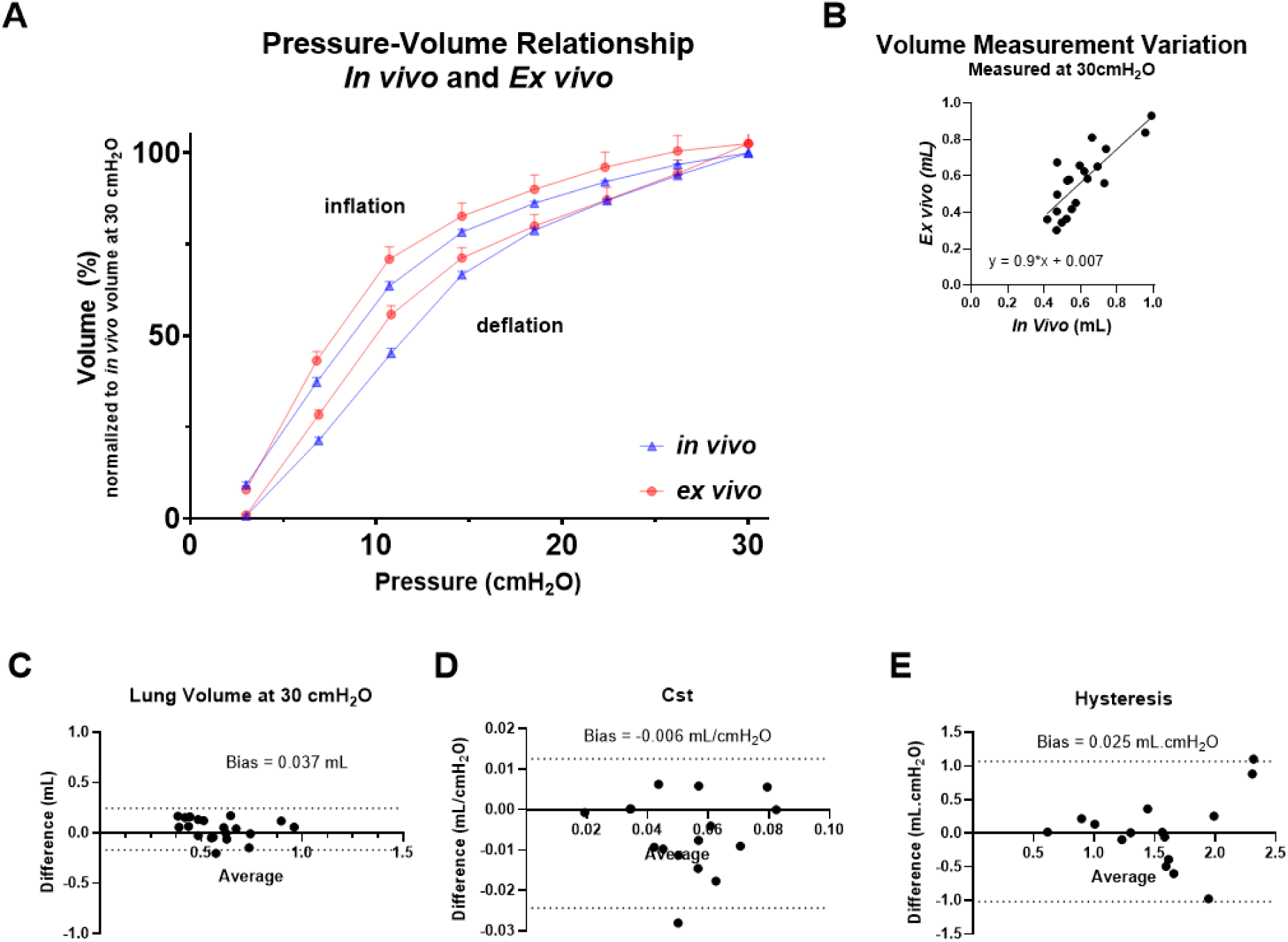
There are no meaningful differences between quasi-static lung function parameters when measured *in vivo* compared to *ex vivo*. A) Mouse specific, pressure-volume loop expressed as a percentage of the ex-vivo volume at 30 cmH_2_O of pressure, inflation and deflation. There is little absolute difference between maximal and minimum volumes lungs are inflation by positive pressure when measured either *in vivo* or *ex vivo*. Mid-pressure volumes are slightly increased when measured *ex vivo*. B) The relationship between volumes due to pressure of lungs are similar when measured *in vivo* or *ex vivo*. C) The difference in lung volume at 30 cmH_2_O of pressure is centered around zero, indicating similar lung volumes at this pressure D) A small negative bias is observed, indicating a small decrease in static compliance *in vivo* E) The difference in hysteresis of the pressure-volume loop is well centered around zero, indicating little systemic bias in measurement *in vivo* as compared to *ex vivo*.

We compared *in vivo* and *ex vivo* measurements of maximum lung volume measured at 30 cmH_2_O from the PFT and found that they correlate (p < 0.05, Pearson r=0.81, 95% CI [0.5908 to 1.2570], n=20, Figure 3B). There is strong agreement between these maximum lung volumes when measured *in vivo* or *ex vivo* (Figure 3C). We found that there is a bias of 37 uL between *in vivo* and *ex vivo* measurements with the limits of agreement being approximately ± 200 uL. This bias is minimal compared to the 300 uL decrease in lung volume seen in mouse models (C57BL6 and BALB/c) of fibrosis (18) and the 300 uL increase as seen in mouse models (Fib5KO) of emphysema (19). Additionally, we found little bias between *in vivo* and *ex vivo* static compliance (Cst) measurements, with a bias of 0.006 mL/cmH_2_O (Figure 3D) and limits of agreement from - 0.024 to 0.013 mL/cmH_2_O. This is again a lower bias compared to other pathological literature values where up to a 0.03 mL/cmH_2_O increase is seen in models of emphysema, and a decrease of similar magnitude is seen in bleomycin models (19, 20). Similarly, we see little bias in hysteresis, the area between the pressure-volume curve (0.025 cm^2^, Figure 3E) with limits of agreement of –1.02 to 1.07. This is much smaller than a 2 cm^2^ increase seen in bleomycin mouse models (18).

Next, we compared *in vivo* and *ex vivo* results for dynamic regime parameters. The limits of agreement (from 0.35 to 5.9 cmH_2_O/mL) for tissue damping, G, are less than physiological meaningful changes (for example, there is ∼7 cmH_2_O/mL increase seen in bleomycin models (18)). However, we found that *ex vivo* measurements are biased by 3.14 cmH_2_O/mL (Figure 4A). Tissue elastance parameter, H shows a slight bias of 2.3 cmH_2_O/mL (Figure 4B), but the limits of agreement appear relatively well-centered around zero, from –14.88 to 19.43 cmH_2_O/mL. It is noted that the bleomycin injury model results in a 40 cmH_2_O/mL increase in H (18). The relatively large change in G compared to that in H causes a downstream bias of 0.066 in hysterisivity, η, calculated as the ratio of G/H (Figure 4C). The limits of agreement were between 0.03 to 0.10. This is meaningful considering bleomycin models of C57Bl/6 mice have shown no change in hysterisivity 21 days post-exposure (18). Newtonian resistance, Rn, showed a bias of 0.03 cmH_2_O.s/mL (Figure 4D) compared to a nearly 0.15 cmH_2_O.s/mL increase seen in cigarette smoke-induced mouse models of COPD (21). Similar to the static compliance, the dynamic compliance, Crs, indicated a mild bias of –0.005 mL/cmH_2_O (Figure 3E), with limits of agreement between –0.019 to 0.008. The bias is less than that seen in bleomycin fibrosis mouse models, of up to a 0.015 decrease in dynamic compliance (20). The parameters with the greatest change are tissue damping and hysterisivity, indicating that the lung is more mechanically heterogeneous *in vivo* compared to *ex vivo*.

**Figure 4.**
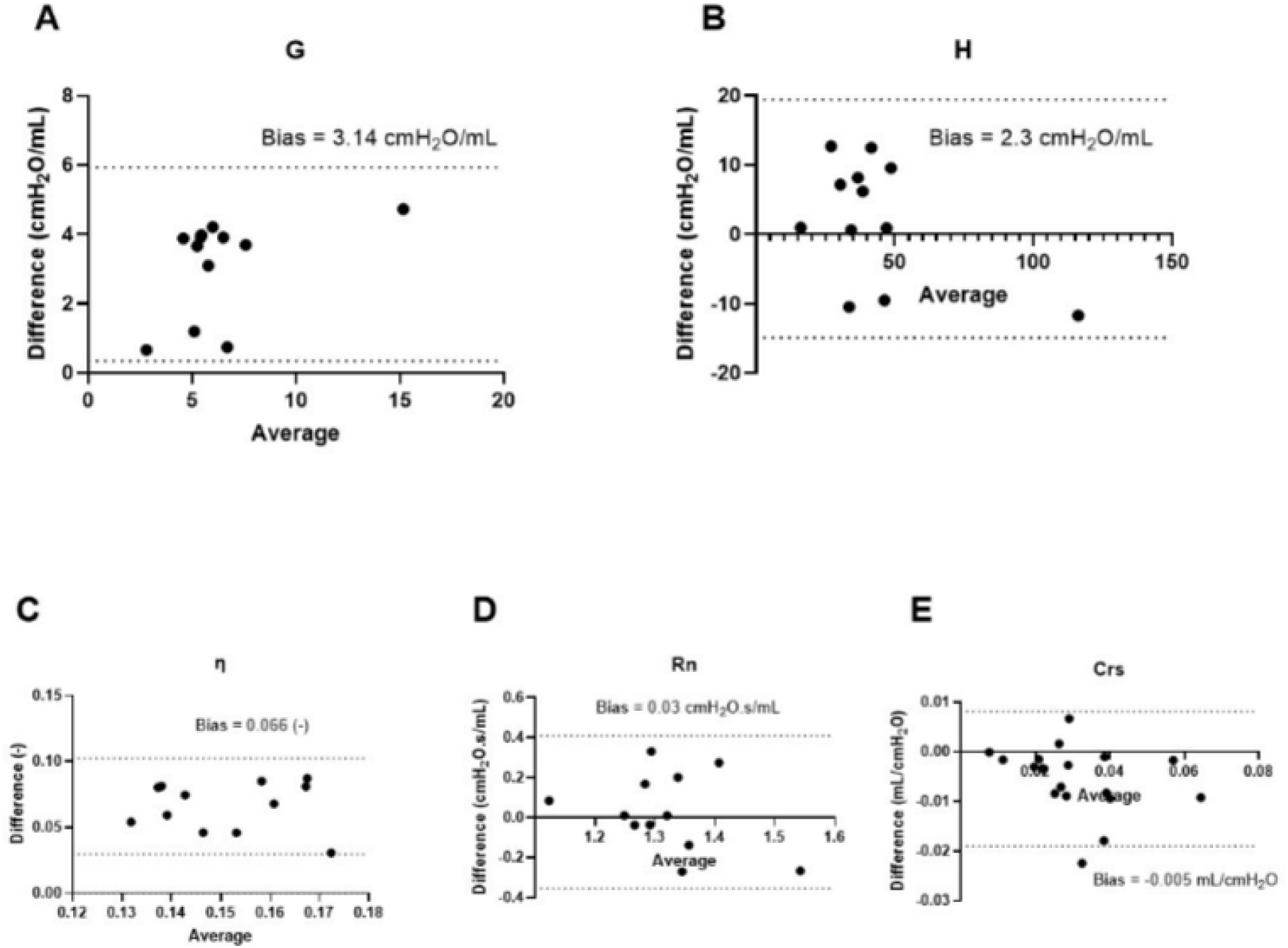
There are meaningful differences in dynamic lung function parameter when measured *in vivo* compared to *ex vivo*. A) Apparent tissue damping increases when measured *in vivo* compared to *ex vivo* B) Hysteresivity, often used as a measure of heterogeneity, increases *in vivo* compared to *ex vivo* C) A smaller positive bias is observed in tissue elastance, indicating a small increase *in vivo* compared to *ex vivo* D) The difference in Newtonian resistance, representative of conducting airway resistance, appears well centered around zero, indicating little systemic bias in its measurement *in vivo* compared to *ex vivo* E) A moderate negative bias is observed in compliance of the respiratory system (dynamic compliance), indicating decreased compliance *in vivo* compared to *ex vivo*.

We performed *u*CT imaging on a subset of the samples (5 male, 3 female) both *in vivo* and *ex vivo* to validate the volumetric findings of the PFTs. From *ex vivo scans*, we segmented whole lung volume. Because the left and accessory lung lobes were commonly not visible *in vivo*, only the right lung lobe volumes were segmented. Comparatively, due to the frozen nature of the *ex vivo* lungs, there was no reliable way to isolate the right lung from the whole lung volume. Thus, we compared the *in vivo* right lung volume to the *ex vivo* whole lung volume. We found a significant relationship between the *ex vivo* volume and the age of the mouse samples (p<0.05); however, no statistically significant relationship between *in vivo* right lung volumes and age (Figure 5A). In directly comparing the *ex vivo* whole lung volume and the *in vivo* right lung volume, no significant relationship was shown (Figure 5B), though a general positive trend was found. One prespecified outlier (exceeding 2 standard deviations from expected values) was excluded. In comparing the *ex vivo* whole lung volume and the *in vivo* whole lung 2D thoracic area (Figure 5C), again no significant relationship and a general positive trend was found. Finally, at 20 cmH_2_O of pressure, lung volume obtained via *ex vivo u*CT imaging was found to have a strong correlation to volumes obtained via *ex vivo* PFT at matched pressures (p<0.05, r=0.83, Figure 5D). At end expiration during normal tidal breathing, the pleural pressure is accepted as being –5 cmH_2_O (22). In this state, the alveoli are at equilibrium with the atmosphere, resulting in an airway opening pressure of 0 cmH_2_O. The transpulmonary pressure, being the difference between airway and pleural pressures is therefore 5 cmH_2_O. During the *ex vivo* PFT, the pleural pressure is 0 cmH_2_O at all times, meaning the transpulmonary pressure is equal to the airway opening pressure. Therefore, the lungs *in vivo* at end expiration during normal tidal breathing should be equivalently loaded to the lungs held at 5 cmH_2_O of airway opening pressure during the *ex vivo* PFT. The volume of the lungs obtained via *u*CT at end expiration was found to have a weak correlation to total lung volume obtained via *ex vivo* PFT at 5 cmH_2_O (Figure 5D).

**Figure 5.**
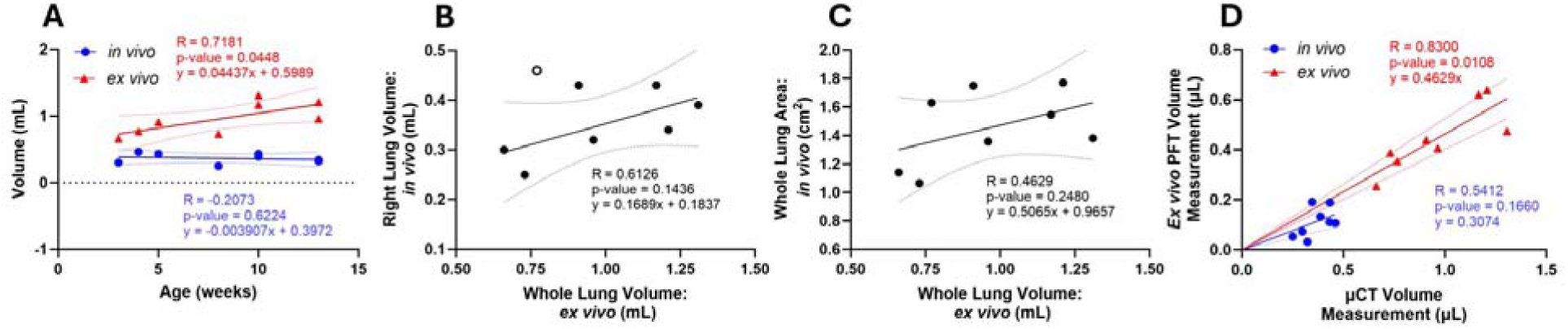
*Ex vivo* imaging at fixed pressures provides better mechanical characterization of lung mechanics than *in vivo* imaging. (A) *In vivo* and *ex vivo* imaging differed in their ability to detect age-related changes in lung volume, highlighting a limitation of *in vivo* imaging for assessing mouse lung mechanical properties. (B) Even when restricting the *in vivo* analysis to the right lung, which appeared well aerated and was easier to segment, correlation with *ex vivo µ*CT lung volume remained poor. (C) Similarly, estimating lung volume by summing 2D axial areas showed poor agreement with *ex vivo* volume measurements. (D) *Ex vivo* µCT volumes at supraphysiologic pressure agreed more closely with PFT volumes (r = 0.83, p = 0.01) than did *in vivo µ*CT volumes at physiologic pressure (r = 0.54, p = 0.17), likely because the higher distending pressure expands the lung toward a more uniform, fully recruited state, reducing the regional variability in aeration that degrades volume measurements.

## DISCUSSION

We hypothesized that thoracic structures (bone, skeletal muscle, diaphragm, and mediastinal anatomical components) influence biomechanical properties. To dissect the underlying mechanical contributions of the chest wall and surrounding structures, we analyzed the PFT results from the quasi-static and dynamic regimes separately. This distinction is informative because the two regimes probe different aspects of lung mechanics: quasi-static measurements reflect near-equilibrium elastic behavior, whereas dynamic measurements capture frequency-dependent properties and effects of regional mechanical heterogeneity.

After analyzing both regimes, the results partially support our hypothesis of thoracic structure influence. Quasi-static pressure-volume behavior such as lung volumes, airway resistance, and static and dynamic compliance were similar when measured *in vivo* and *ex vivo*. In contrast, tissue damping (G) and hysterisivity (η) differed substantially between conditions. We infer that the parameters that vary across *in vivo* and *ex vivo* conditions are influenced by thoracic structures extrinsic to the lung. As such, these thoracic structures contribute relatively little to elastic properties of lung parenchyma but substantially to apparent energy dissipation. Collectively, these findings suggest that several, but not all, clinically relevant mechanical properties of the lung are intrinsic to airway and parenchymal tissue.

### Static Regime Parameters

Pressure-volume behavior in the quasi-static regime was preserved between *in vivo* and *ex vivo* measurements. This indicates that maximal lung inflation, quasi-static compliance, and quasi-static elasticity are intrinsic homeostatic biomechanical setpoints of the airways and parenchymal tissue rather than a result of restriction from surrounding thoracic structures. The main differences between *in vivo* and *ex vivo* measurements emerged at lower pressures during inflation. Because the differences only presented at lower pressures, we infer that the thoracic boundary conditions primarily influence recruitment and decruitment of alveoli, rather than altering intrinsic elastic recoil of parenchyma. This recruitment modulation role is consistent with clinical literature showing that external loads associated with obesity, abdominal distension, and supine posture promote dependent atelectasis and ventilation heterogeneity (23–26).

These conclusions are supported by similar high-pressure deflation behavior, the strong correlation and agreement of lung volume measurements at 30 cmH_2_O, and the small biases in static compliance and PV-loop hysteresis. Together, the quasi-static data suggest that thoracic structures primarily contribute to the shift in pressure ranges over which lung alveoli open and close with relatively little contribution to elasticity of lung parenchyma under near-equilibrium conditions.

### Dynamic Regime Parameters

In the dynamic regime, elastic and conductive behaviors were largely preserved between *in vivo* and *ex vivo* conditions, with the clearest differences reflecting an increased apparent dissipation *in vivo*. Central airway resistance (Rn) showed little bias, arguing against changes in proximal airway caliber as the primary driver. Likewise, dynamic measures of elasticity were only modestly different. The constant phase model’s tissue elastance parameter (H) saw a small increase *in vivo*, but this increase is inherent to H as the parameter incorporates any residual external elastic load coupled to the parenchyma and cannot fully separate intrinsic parenchymal stiffness from thoracic boundary conditions. Together with the small changes in static and dynamic compliance, these findings indicate that thoracic structures external to the lung have limited impact on elastic stored energy.

By contrast, tissue damping (G) shows a clear *in vivo* increase, driving a corresponding increase in hysterisivity (η = G/H). Notably, there was little change in quasi static hysteresis, suggesting that the added dissipation occurs primarily under dynamic and frequency dependent conditions. One plausible explanation is the increased biomechanical heterogeneity *in vivo*, which promotes asynchronous filling and redistribution of air (as in pendelluft) due to original time constant mismatches (27). This process inflates tissue damping (G, the real component of impedance) in the constant phase model without requiring large changes in intrinsic parenchymal viscoelasticity (H, the imaginary component of impedance) (28). We’ve proposed that a non-uniform thoracic loading of the rib cage and diaphragm accentuates regional differences in compliance that diminish *ex vivo*. However, we acknowledge that heterogeneous regional airway caliber could contribute similarly, which is not ruled out via a nearly unchanged Newtonian resistance (Rn).

There is limited information regarding regional biomechanical heterogeneity of intrinsic properties such as compliance. This warrants future investigation. Our findings align with prior work showing that heterogeneity can overestimate tissue damping (G), and with studies reporting higher hysterisivity (η) in intact or open-chest conditions compared to isolated lungs, in which reduced heterogeneity in the isolated lung shifts η toward values measured in tissue strips (29).

### Physiological context: comparison to disease models

To contextualize the magnitude of thoracic-structure effects, we compare our *in vivo* and *ex vivo* differences to established murine disease models. This frame of reference helps determine whether the modest changes we observe in elastic and conductive metrics are physiologically meaningful and clarifies which mechanical signatures are more consistent with thoracic boundary conditions versus parenchymal pathology.

Relative to bleomycin-induced fibrosis, thoracic structures produced a smaller and qualitatively different pattern of change. Our *ex vivo* measurements saw damping increased substantially (∼52%) while elastance increased only modestly (∼9%), whereas fibrosis increases both damping (∼200%) and elastance (∼100%) markedly as collagen accumulates (18). Mechanistically, this distinction is informative: fibrotic remodeling stiffens the tissue (raising H) and increases dissipation (raising G) in a way that tends to preserve η, consistent with structural damping of altered parenchyma. The preferential elevation of G (and η) with comparatively preserved H is more consistent with added mechanical heterogeneity than with a primary change in intrinsic tissue material properties. Concordantly, bleomycin injury also produces prominent PV-curve changes, hysteresis changes, and a reduced volume at 30 cmH_2_O, whereas thoracic structures have negligible effects on those quasi-static indices (18). There is a similar contrast for compliance: thoracic structures reduce compliance modestly (∼11%), whereas bleomycin produces a substantially larger decrease (∼40%) (20).

Models of emphysema provide a complementary comparison. In murine models of emphysema, dynamic compliance increases markedly (∼42%) and Newtonian resistance rises (∼19%), reflecting loss of elastic recoil and airway instability (30). In our data from *ex vivo* measurements, thoracic structures shift dynamic compliance modestly in the opposite direction (-17%) and leave large airway resistance largely unchanged (2%). Both of these components indicate that *ex vivo* mechanics do not recapitulate either restrictive (fibrosis) or obstructive (emphysema) disease signatures. Instead, the thoracic boundary condition’s distinctive fingerprint is a selective elevation of dynamic dissipation (G, η) with relatively preserved elastic storage and conductive airway properties. These comparisons support the interpretation that the thoracic contribution to lung biomechanics is real but sub-pathological in magnitude.

### Volumetric Analysis from **μ**CT imaging

The *in vivo* and *ex vivo* volumetric analysis revealed inconsistencies in the lung volumes derived from their respective imaging modalities. *In vivo* volumetric analysis had no significant or positive trend between lung volume and age. This was surprising, as murine lung volumes increase with age across the range of our mouse models (3 to 13 weeks) in both male and female mice. A potential contributor to these results was the obstruction of left and accessory lobes in our μCT imaging. However, when attempting to correct for this with thoracic area measurements, there was still no trend found. For the *ex vivo* images, we found a positive, statistically significant relationship between age and lung volume. To verify our measurement approaches, we compared the volumes measured from both the μCT imaging (*in vivo* and *ex vivo*) volumes to their respective PFT volumes at comparable loading conditions. We found the *ex vivo* methods had a statistically significant correlation, while *in vivo* samples showed only a positive trend. Given the added difficulty of live animal imaging and the inconsistencies for *in vivo* volume quantifications, *ex vivo* imaging may offer a more straightforward method for quantifying lung morphometry.

In considering differences between imaging techniques, loading also becomes an important factor. *Ex vivo* samples were analyzed at elevated pressures compared to those present during free-breathing, as experienced by the *in vivo* samples. Indeed, our murine PFTs also examined pressure ranges more commonly experienced during mechanical ventilation than during physiological activity. Thus, differences within the smaller loading regime *in vivo* may have been muted, leading to more difficulty in distinguishing the expected volumetric effects of maturation. In contrast, *ex vivo* loading allowed for accentuation of the differences in volume and good agreement with matched load volumes from *ex vivo* PFTs. Together, these effects suggest that supraphysiologic loading *ex vivo* could allow for better morphometric characterization of lung differences.

### Synthesis and implications

From this experiment, we conclude that the chest wall and surrounding structures exert both a dissipative and a mildly restrictive effect on the respiratory system, evidenced by the increase in tissue damping and the decrease in compliance *in vivo*. We further conclude that these effects are not uniformly distributed throughout the lung but are heterogeneous, accounting for the increase in hysterisivity *in vivo*.

The nearly identical pressure-volume relationship at higher inflations suggests that, in the mature lung, maximal expansion is set largely by intrinsic parenchymal recoil rather than acute chest-wall constraint. In a developmental context, where mechanical stretch guides growth and remodeling and compensatory capacity declines with age (31, 32), this provides biomechanical insights to the growing body of literature that mechanical changes to the lung during early life impacts adult parenchymal properties (33).

Clinically, the central implication is that abnormal chest-wall contributions may present more strongly in dissipation- and heterogeneity-sensitive metrics (G, η) than in compliance, potentially rendering dysfunction less apparent to conventional compliance-based monitoring. This framework is directly relevant to situations where thoracic loading changes abruptly (e.g., open-chest surgery) and to conditions imposing nonuniform external loads (e.g., obesity and other chest-wall/diaphragm or pleural disorders) (24, 34–39). It also points to a measurement opportunity: oscillometry-based methods, which are inherently sensitive to frequency-dependent heterogeneity, may better capture chest-wall–driven dysfunction than volume- or compliance-only testing. Finally, because heterogeneity concentrates regional stress and is linked to VILI/ARDS risk (40) interventions that redistribute external loading—especially patient positioning—may help homogenize ventilation and reduce injurious stress concentrations.

### Limitations

These measurements are global, so we cannot localize the source of heterogeneity: the proposed mechanism of nonuniform external loading creating regional time-constant mismatch is inferred rather than directly observed, and our lumped Newtonian resistance does not rule out a contribution from heterogeneous regional airway caliber (either could inflate apparent G). The *in vivo*-to-*ex vivo* sequence could not be randomized, so recruitment history and loss of perfusion (with potential time-dependent surfactant effects) may partly influence low-pressure PV behavior, though these factors are unlikely to explain the elevated dynamic dissipation at the operating point. Our cohort’s age range increases variance in absolute mechanics, but the paired within-animal design mitigates this for the primary comparisons. Finally, results are bounded by species/posture (anesthetized, supine, ventilated mice) and by use of the constant-phase model, whose homogeneity assumption means *in vivo* parameters should be interpreted as effective, model-derived values rather than literal intrinsic tissue properties

## Supplemental Information

At time of peer-reviewed publication, supplemental data will be available.

Supplemental Video 1: Deep Inflation Maneuver

Supplemental Video 2: Single Frequency Forced Oscillation Technique

Supplemental Video 3: Broadband Forced Oscillation Technique

Supplemental Video 4: Pressure-Volume Loop

## Funding and acknowledgements

This work was supported by the VA VISN1 Fred Wright CDA1, National Institute on Aging R03AG074063, and EPM is a Pepper Scholar of the Yale Claude D. Pepper Older Americans Independent Center supported by NIA P30AG021342. JMS is a Parker B Francis Fellow. MSG is supported by NIH T32 HL007778.

The authors acknowledge the use of the Materials Characterization Facility at Carnegie Mellon University supported by grant MCF-677785. A portion of this research was sponsored by the Army Research Laboratory and was accomplished under Cooperative Agreement Number W911NF-20-2-0175. The views and conclusions contained in this document are those of the authors and should not be interpreted as representing the official policies, either expressed or implied, of the Army Research Laboratory or the U.S. Government. The U.S. Government is authorized to reproduce and distribute reprints for Government purposes notwithstanding any copyright notation herein.

This research was conducted using the ORCHARD cluster. The authors acknowledge Carnegie Mellon University for making this shared high-performance computing resource available to the CMU research community.

